## Supplementary Figures for "Multistep loading of PCNA onto DNA by RFC"

### Supplemental Figures

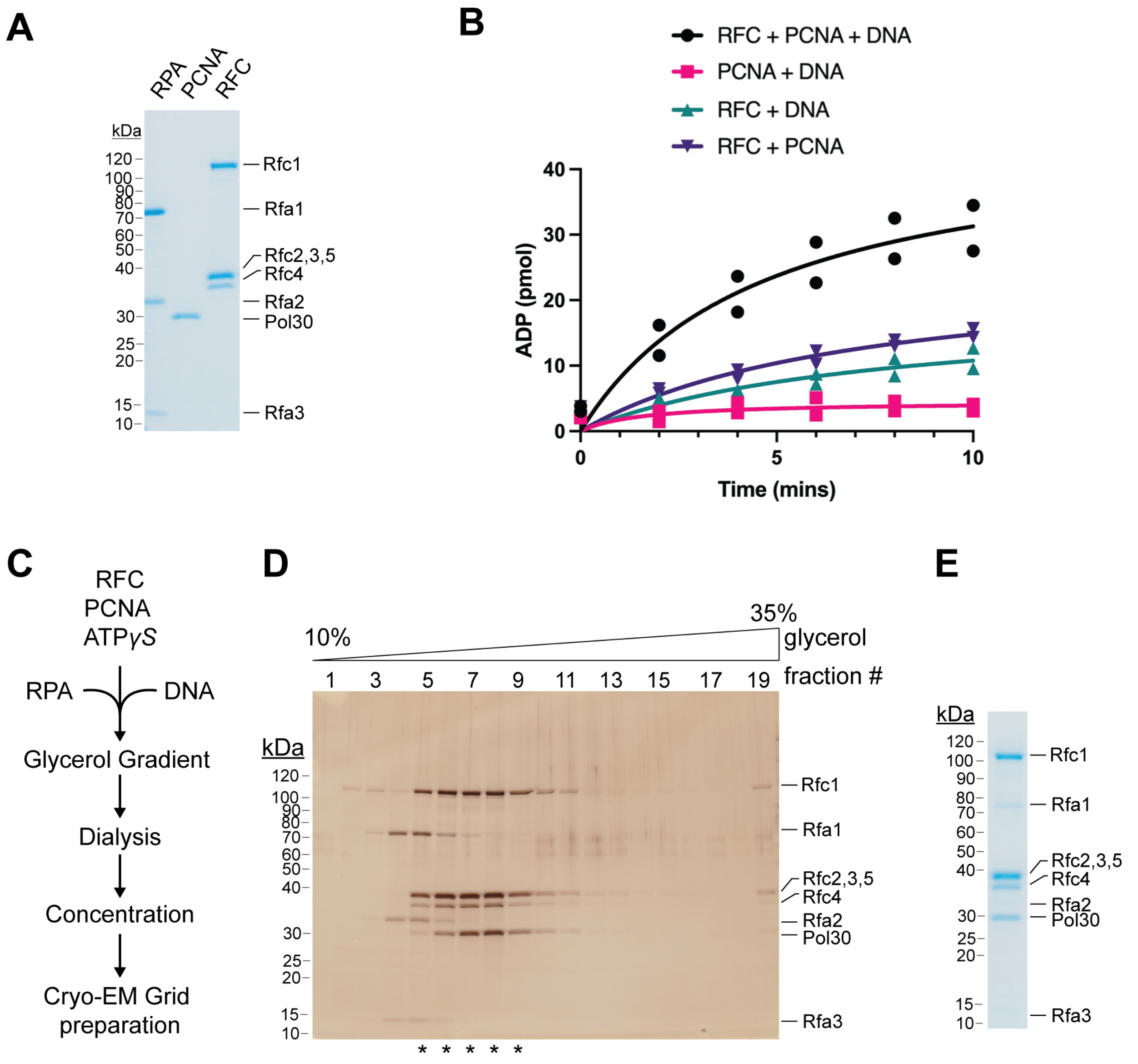


**Figure S1. Purification and analysis of RFC:PCNA. A**, Representative Coomassie-stained SDS-PAGE analysis of purified *S. cerevisiae* RFC, PCNA and RPA. **B**, ATPase activity of RFC in the presence of PCNA and DNA (black circles), RFC in the presence of DNA (teal triangles), RFC in the presence of PCNA (purple triangles) and PCNA and DNA (pink squares). Experiments are shown in duplicate. **C**, Schematic for the assembly of RFC:PCNA:DNA in the presence of ATPγS. **D**, Representative silver-stained SDS-PAGE analysis of RFC:PCNA fractions following glycerol gradient (10-35%) centrifugation. Fractions pooled for Cryo-EM analysis are denoted with *. **E**, Representative Coomassie-stained SDS-PAGE analysis of purified *S. cerevisiae* RFC:PCNA:DNA.

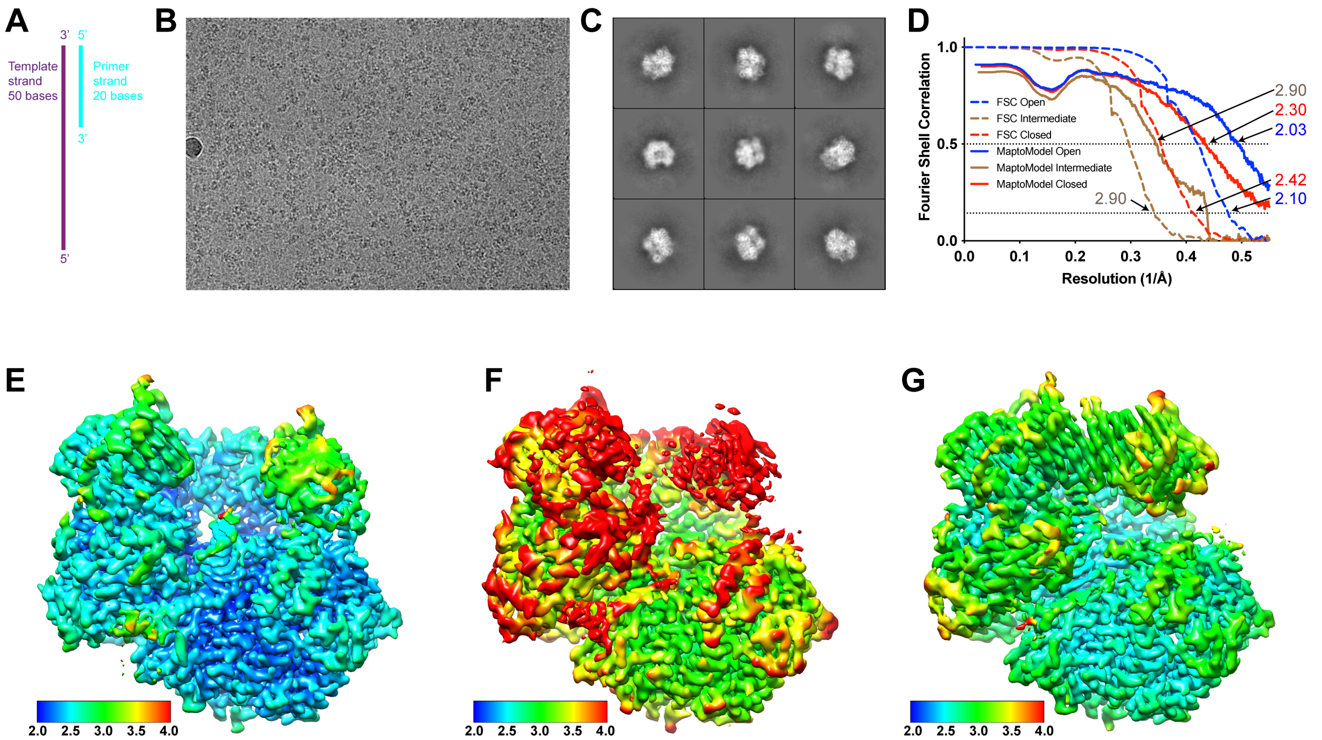


**Figure S2. Cryo-EM analysis of Rfc-RFC:PCNA with DNA substrate 1. A,** Schematic of DNA substrate 1 ($DNA_1$). **B,** Representative cryo-EM image of vitrified RFC:PCNA:$DNA_1$. **C,** Representative two-dimensional averages of RFC:PCNA:$DNA_1$. **D,** Plot of Fourier shell correlations between two independent open state half-maps (dashed blue), two independent intermediate state half-maps (dashed brown), two independent closed state half-maps (dashed red), the open state map and open state atomic model (blue), the intermediate state map and intermediate state atomic model (brown), and the closed state map and closed state atomic model (red). **E-G,** Cryo-EM density maps of open (E), closed (F) and intermediate (G) maps colored by local resolution in Å.

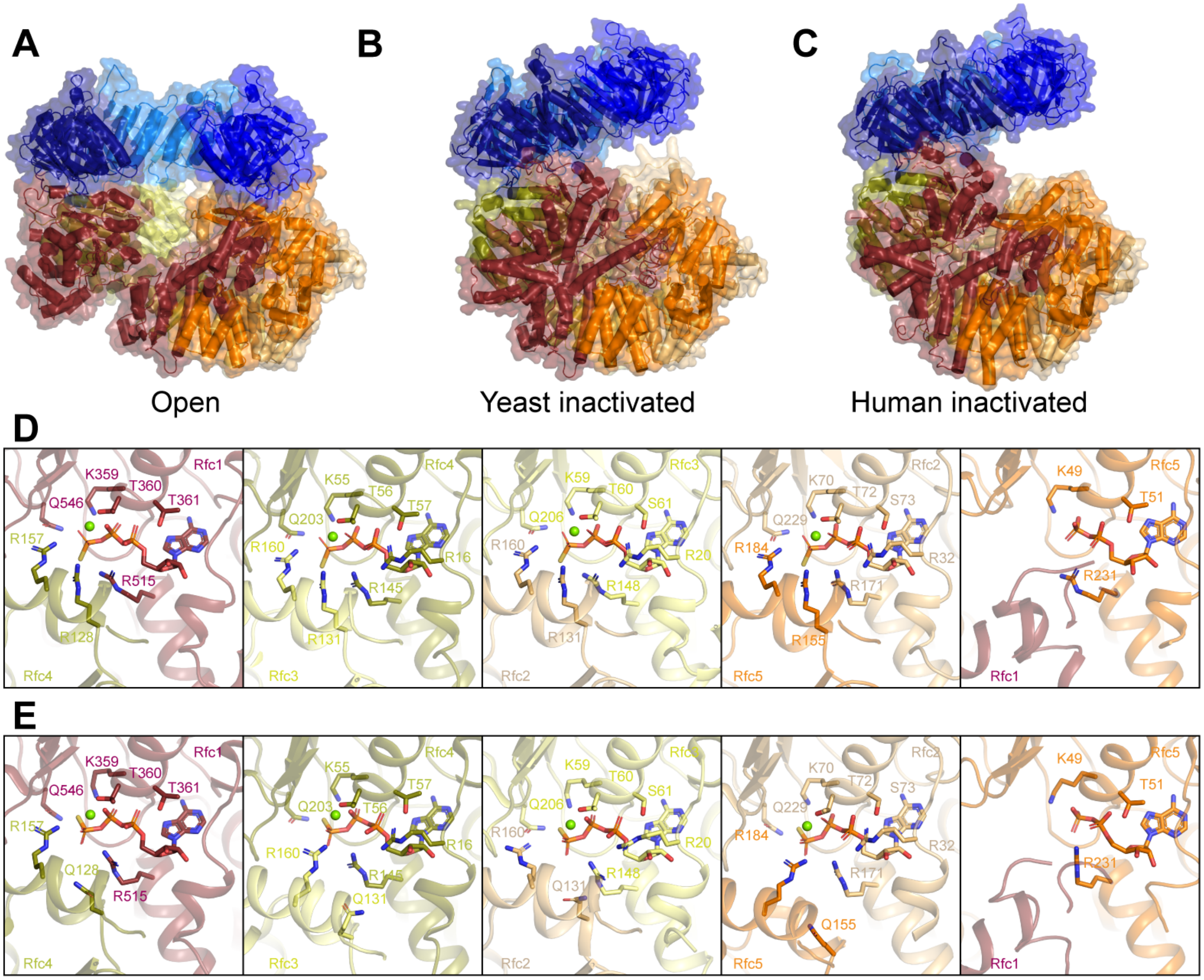


**Figure S3. Comparison of Rfc-RFC:PCNA:DNA$_1$ with structures of yeast and human RFC in inactive states bound to PCNA. A-C,** Structures of open Rfc-RFC:PCNA:DNA$_1$ (A), inactive yeast Rfc-RFC:PCNA (B, PDB: 1SXJ) and inactive human RFC:PCNA (C, PDB: 6VVO). Structures are colored by subunit as in Figure 1. **D-E,** Rfc1, Rfc4, Rfc3, Rfc2 and Rfc5 nucleotide binding sites in Rfc-RFC:PCNA:DNA$_1$ (D) and inactive yeast Rfc-RFC:PCNA (PDB: 1SXJ, E).

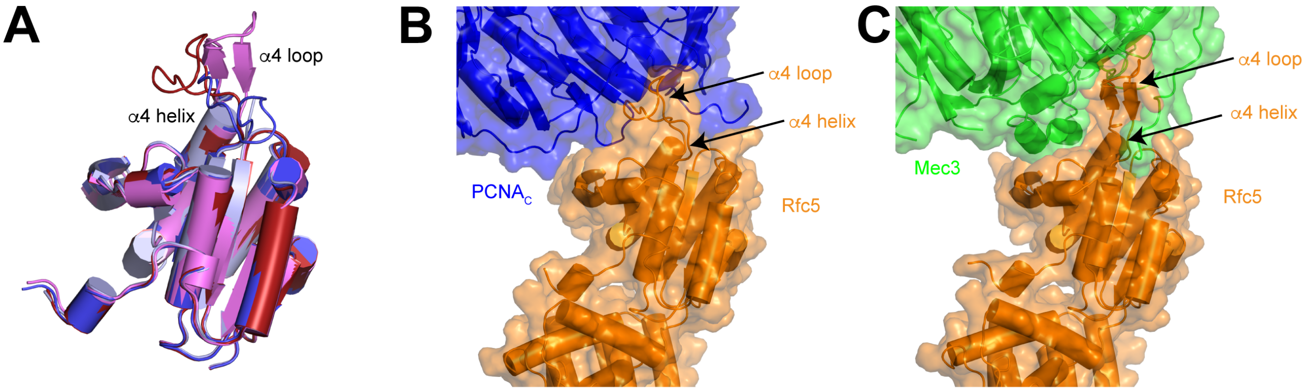


**Figure S4. Conformational flexibility of Rfc5 α4 loop. A,** Interface between Rfc5 and $PCNA_C$ in open state. **B,** Interface between Rfc5 and Mec3 in an open structure of Rad24-RFC in complex with 9-1-1 and DNA (PDB:7ST9). **C,** Superposition of Rfc5 in Rfc-RFC:PCNA:$DNA_1$ open (red), Rfc-RFC:PCNA:$DNA_1$closed (blue) states, Rad24-RFC:9-1-1:DNA (violet) in open and closed states (light blue).

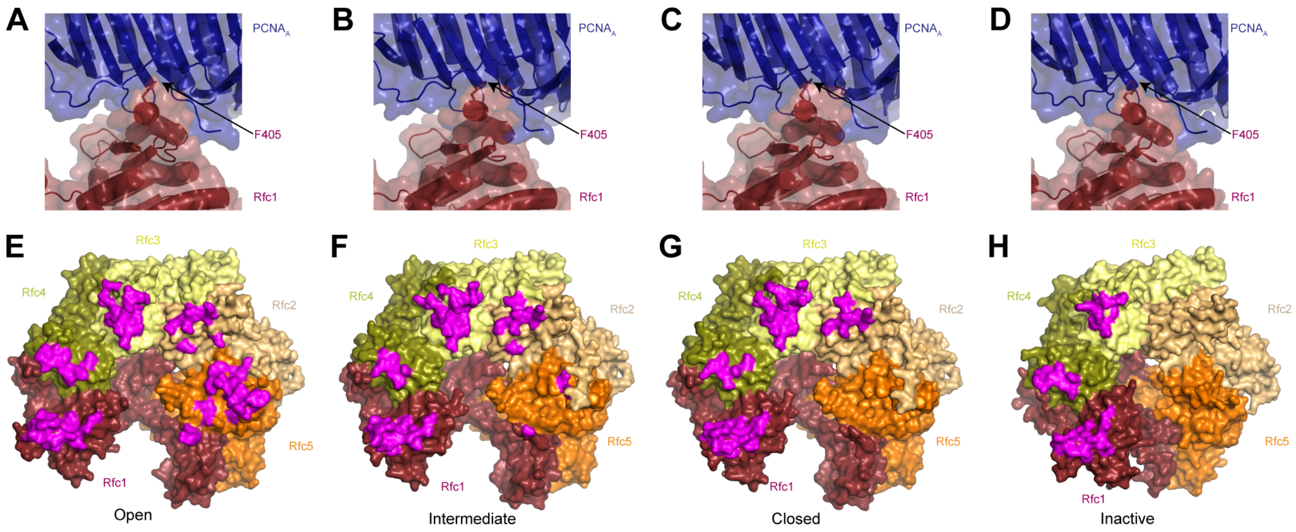


**Figure S5. Rfc1, Rfc4 and Rfc3 form a stable interface with PCNA in active and inactive structures. A-D,** Rfc1-PCNA$_A$ interface in open (A), closed (B), intermediate (C) and inactive (D) states. **E-G,** Structures of RFC in open (E), closed (F), intermediate (G) and inactive (H) states with residues that interact with PCNA highlighted in magenta.

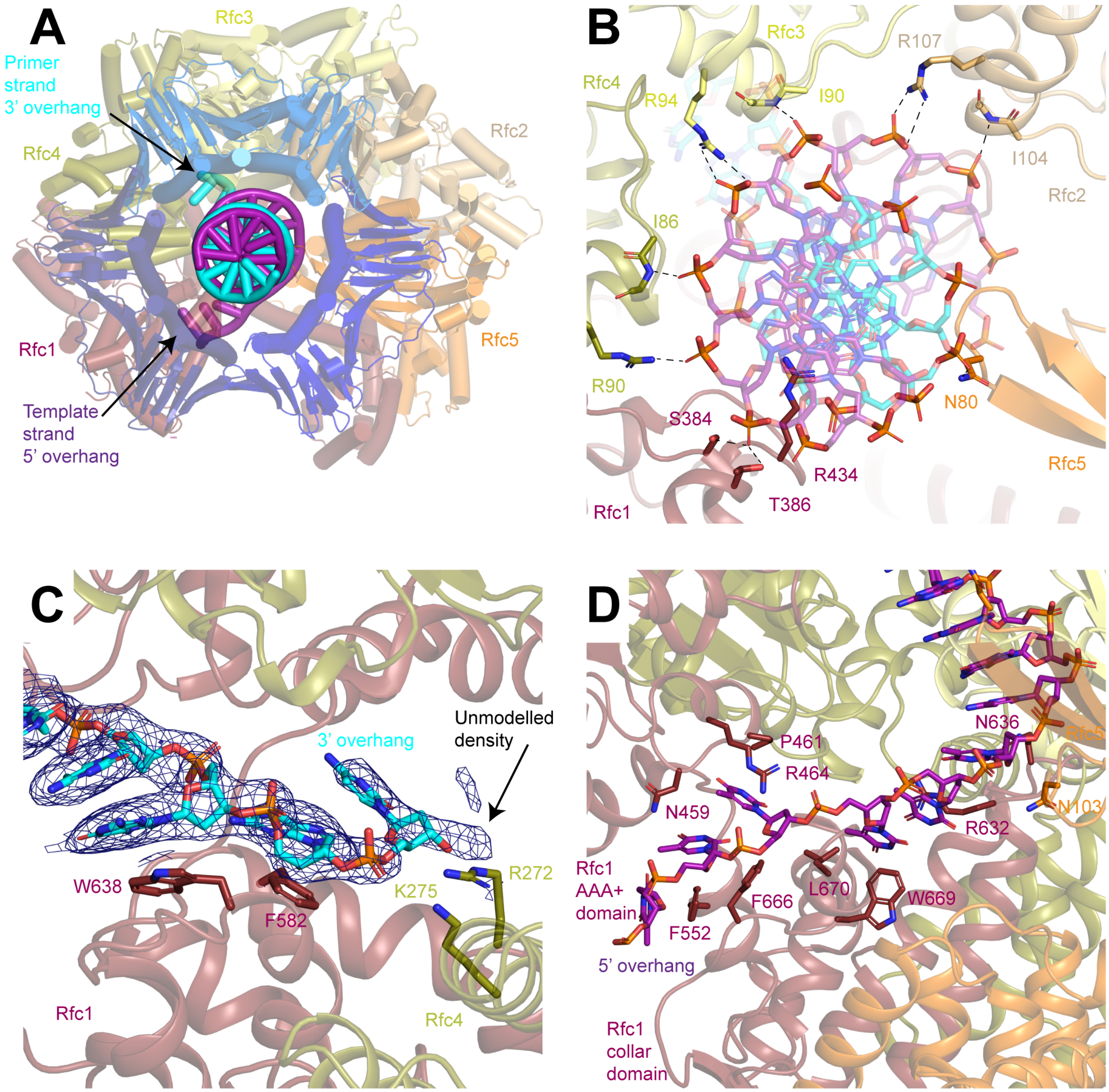


**Figure S6. 3' ss/dsDNA junction binding site 1. A,** Structure of RFC in a closed state with the 3' ss/dsDNA junction at binding site 1, viewed down the central axis of RFC:PCNA. **B,** Coordination of the double-stranded region of the DNA by RFC. Dashed lines represent polar interactions between RFC and the DNA backbone. **C,** Coordination of the 3' overhang of the primer strand by Rfc1 and Rfc4. Density is shown as blue mesh at a threshold of 7 $\sigma$. Arrow points to unmodelled density beyond the last modelled base. **D,** Coordination of the 5' overhang of the template strand by Rfc1.

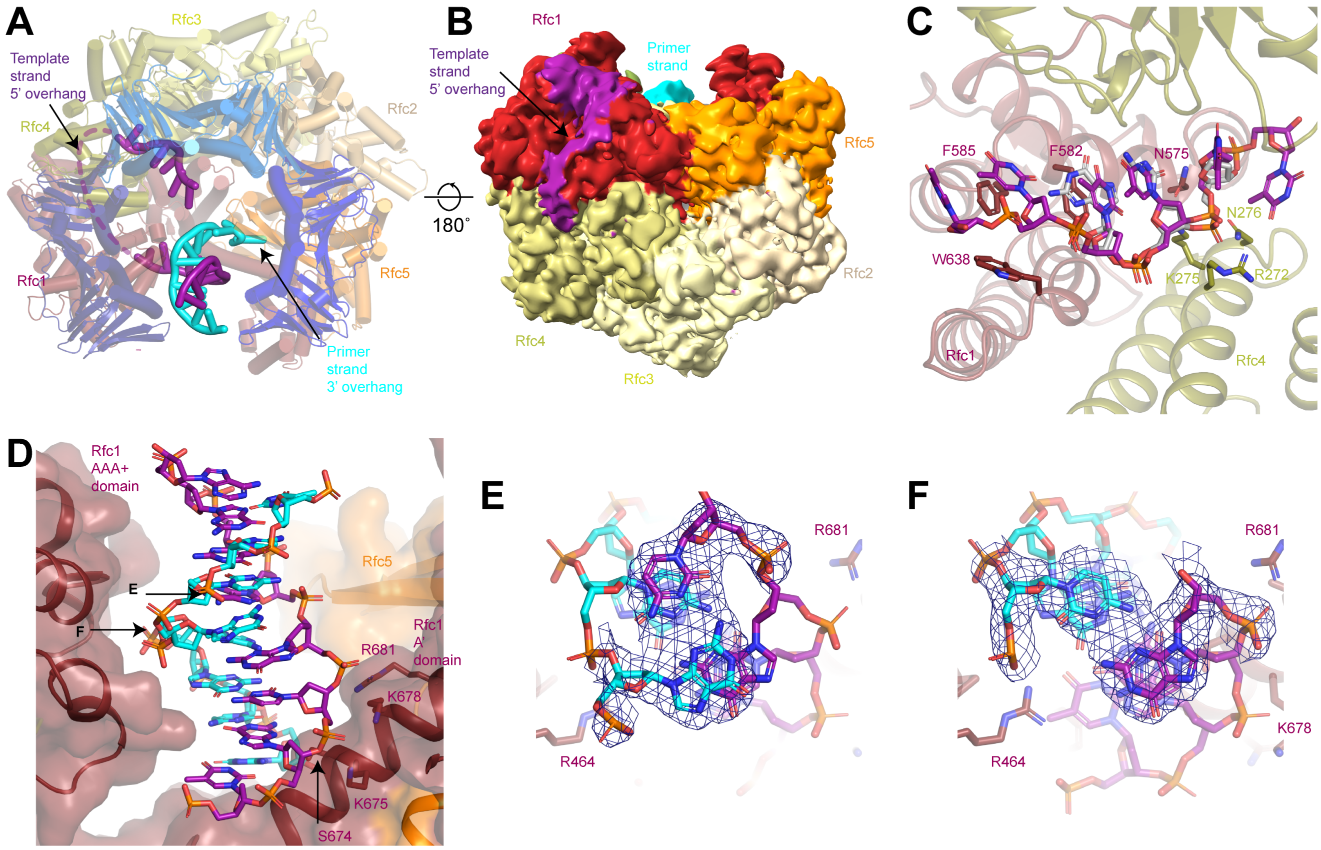


**Figure S7. 3' ss/dsDNA junction binding site 2.** Structure of RFC in an open state with the 3' ss/dsDNA junction at binding site 2, viewed down the central axis of RFC:PCNA. **B,** 5 Å low pass filtered cryo-EM density map of RFC in an open state colored by subunit. Arrow points to poorly ordered density corresponding to the 5' overhang of the template strand. **C,** Coordination of the terminal region of the 5' overhang of the template strand in site 2 (magenta) and the 3' overhang of the primer strand in site 1 (grey) by Rfc1 and Rfc4. **D,** Structure of the double-stranded region of the DNA bound to site 2 in the open state. RFC is shown as a surface. Arrows point to bases that do not adopt Watson-Crick base pairing, which are highlighted in E and F. **E,F,** Density map and model for bases that do not adopt Watson-Crick base pairing. Density is contoured around the highlighted bases and shown as blue mesh at a threshold of 8 $\sigma$.

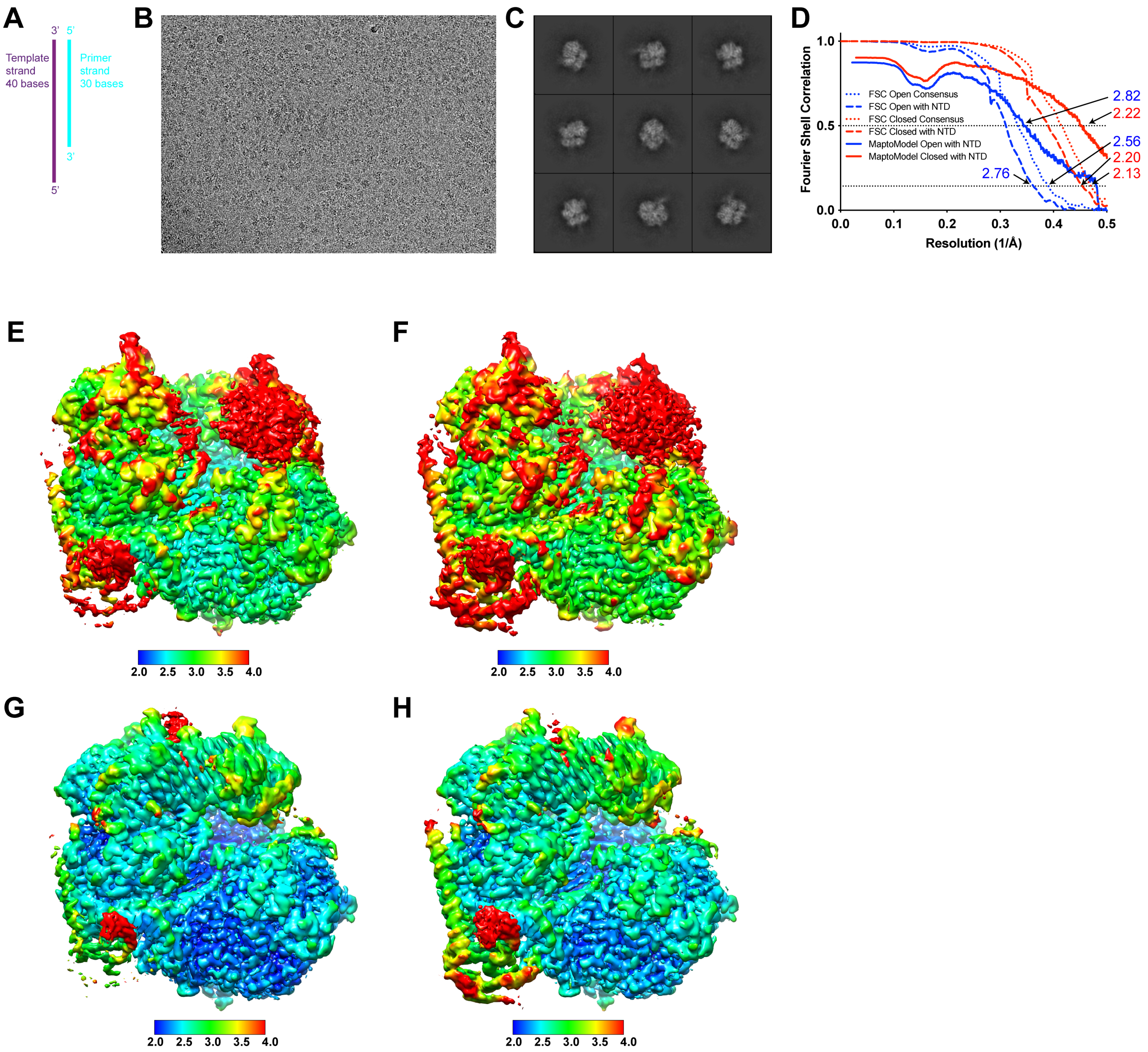


**Figure S8. Cryo-EM analysis of Rfc-RFC:PCNA with DNA substrate 2. A,** Schematic of DNA substrate 2. **B,** Representative cryo-EM image of vitrified RFC:PCNA:$DNA_2$. **C,** Representative two-dimensional averages of RFC:PCNA:$DNA_2$. **D,** Plot of Fourier shell correlations between two independent consensus open state half-maps (dotted blue), two independent open state with NTD half-maps (dashed blue), two independent consensus closed state half-maps (dotted red), two independent closed with NTD state half-maps (dashed red), the open state with NTD map and open state with NTD atomic model (solid blue), the closed state with NTD map and closed state with NTD atomic model (solid red). **E-H,** Cryo-EM density maps of open consensus (E), open with NTD (F), closed consensus (G) and closed with NTD maps colored by local resolution in Å.

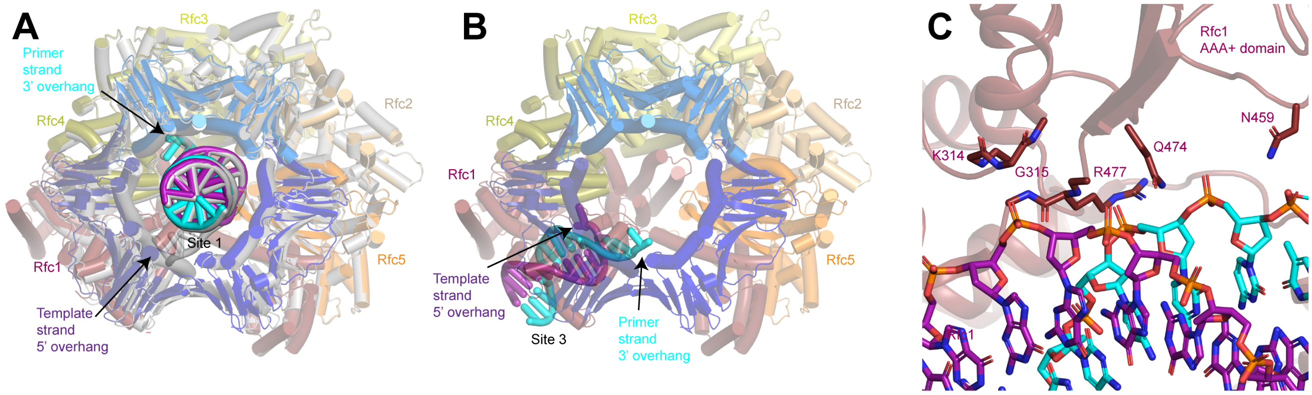


**Figure S9. Coordination of two 3' ss/dsDNA junctions by RFC. A,** Superposition of closed RFC:PCNA with DNA substrate 2 at sites 1 and 3 (color by subunit) and closed RFC:PCNA with DNA substrate 1 at sites 1 (grey). **B,** Structure of RFC in an closed state with the 3' ss/dsDNA junction at binding site 3, viewed down the central axis of RFC:PCNA. **C,** Coordination of the double-stranded region of 3' ss/dsDNA junction at binding site 3 by Rfc1.

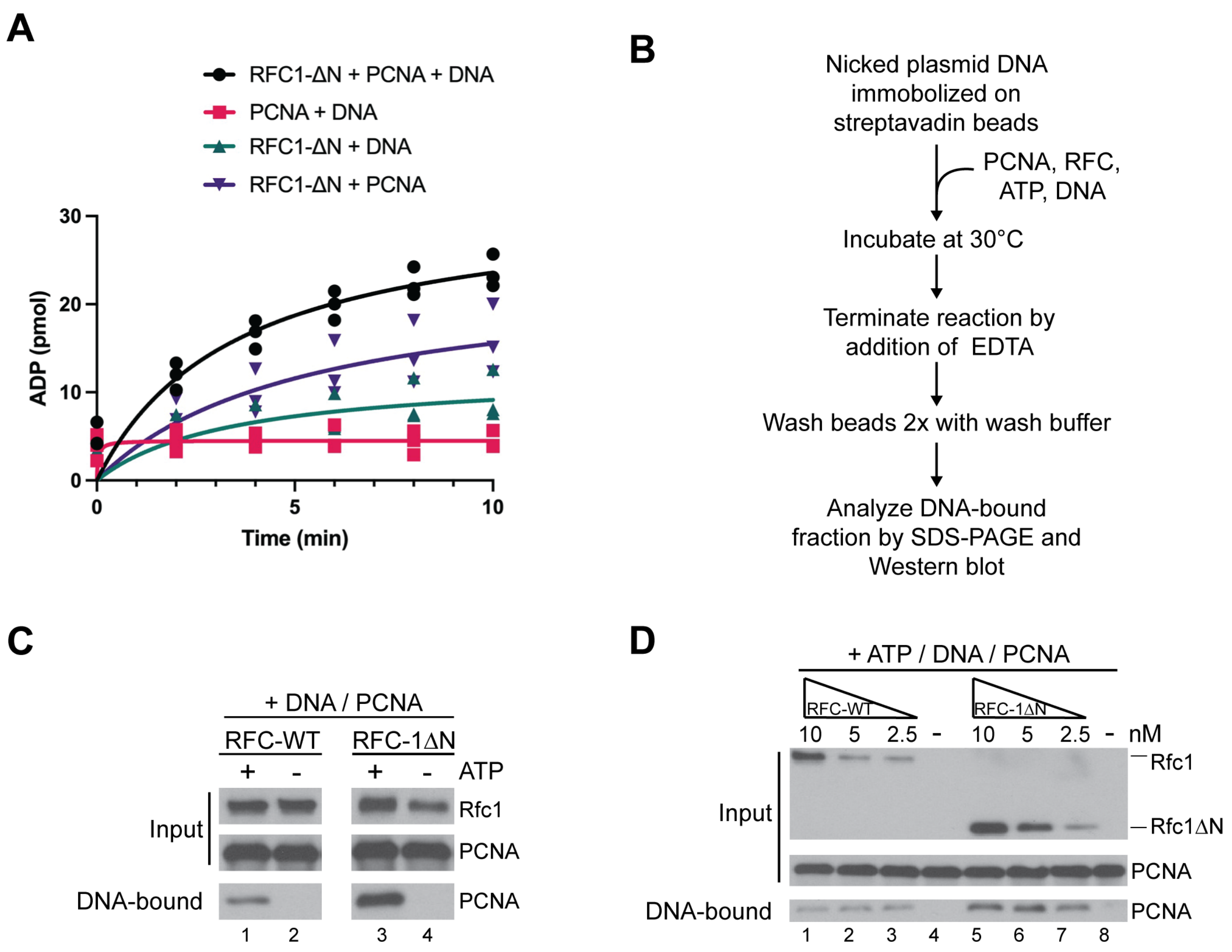


**Figure S10. Analysis of PCNA loading activity of RFC-1ΔN**. **A**, ATPase activity of RFC-1ΔN in the presence of PCNA and DNA (black circles), RFC-1ΔN in the presence of DNA (teal triangles), RFC-1ΔN in the presence of PCNA (purple triangles) and PCNA and DNA (pink squares). Experiments are shown in triplicate. **B**, Schematic for PCNA loading assay using bead-immobilized nicked plasmid DNA as template. **C**, ATP-dependence of PCNA loading activity by RFC-WT (lanes 1 + 2) and RFC-1ΔN (lanes 3 + 4). **D**, Analysis of the dependency of PCNA loading on RFC-WT (lanes 1 - 4) and RFC-1ΔN (lanes 5 - 8). RFC or RFC-1ΔN were included in reactions containing DNA-beads, ATP and PCNA at the concentrations indicated on the top. All experiments were replicated at least two times.

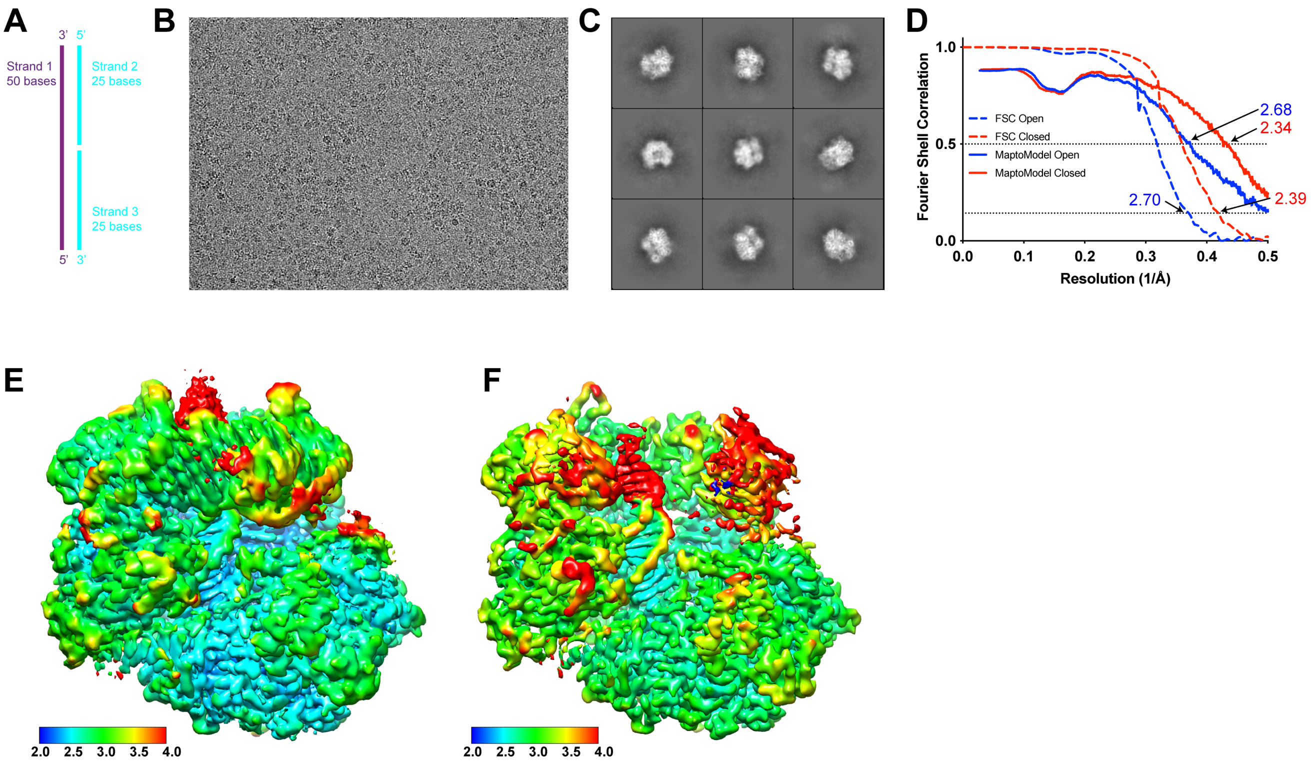


**Figure S11. Cryo-EM analysis of Rfc-RFC:PCNA with a nicked DNA. A,** Schematic of a nicked DNA substrate (nDNA). **B,** Representative cryo-EM image of vitrified RFC:PCNA:nDNA. **C,** Representative two-dimensional averages of RFC:PCNA:nDNA. **D,** Plot of Fourier shell correlations between two independent open state half-maps (dashed blue), two independent closed state half-maps (dashed red), the open state map and open state atomic model (solid blue), and the closed state map and closed state atomic model (solid red). **E-G,** Cryo-EM density maps of closed (**E**) and open intermediate (**F**) maps colored by local resolution in Å.

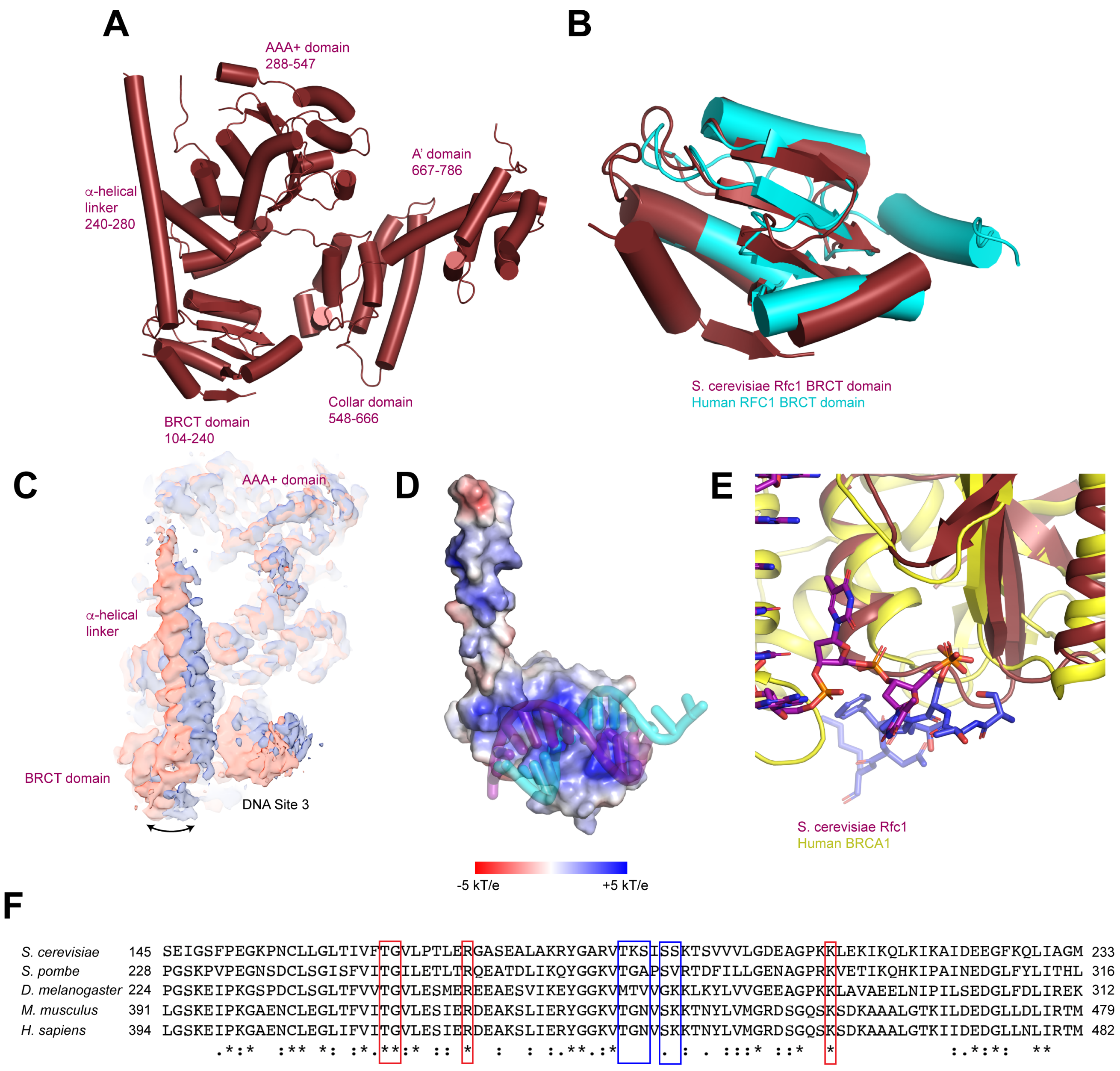


**Figure S12. Structure of the Rfc1 N-terminal domain, A,** Domain organization of Rfc1. **B,** Superposition of the BRCT domain of yeast Rfc1 (maroon) with the BRCT domain of human RFC1 (PDB: 2K6G, cyan). **C,** Cryo-EM density maps for 2 sub-classes of RFC:PCNA with DNA substrate 2 in a closed state. The maps are aligned by the core of RFC. **D,** Rfc1 NTD-DNA interface. Rfc1 NTD is shown as a surface colored by electrostatic surface potential. **E,** Superposition of BRCT domain of yeast Rfc1 (maroon) with the BRCT domain of human BRCA1 in complex with a phosphorylated peptide (PDB:1T15, yellow/blue). **F,** Sequence alignment of the BRCT domains of *S. cerevisiae, S. pombe, D. melanogaster, M. musculus*

and *H. sapiens* Rfc1 orthologues. Residues boxed in red bind to the 5’ overhang and residues boxed in blue bind to the double-stranded region of the DNA.

**Table 1. Cryo-EM data collection, refinement and validation statistics**

| | Rfc1-RFC-DNA1 open (EMDB-xxxx) (PDB xxxx) | Rfc1-RFC-DNA1 intermediate (EMDB-xxxx) (PDB xxxx) | Rfc1-RFC-DNA1 closed (EMDB-xxxx) (PDB xxxx) | Rfc1-RFC-DNA2 closed consensus (EMDB-xxxx) (PDB xxxx) | Rfc1-RFC-DNA2 closed with NTD (EMDB-xxxx) (PDB xxxx) | Rfc1-RFC-DNA2 open consensus (EMDB-xxxx) (PDB xxxx) | Rfc1-RFC-DNA2 open with NTD (EMDB-xxxx) (PDB xxxx) | Rfc1-RFC-nicked DNA closed (EMDB-xxxx) (PDB xxxx) | Rfc1-RFC-nicked DNA open (EMDB-xxxx) (PDB xxxx) |
|---|---|---|---|---|---|---|---|---|---|
| **Data collection and processing** | | | | | | | | | |
| Magnification | 29,000x | 29,000x | 29,000x | 29,000x | 29,000x | 29,000x | 29,000x | 29,000x | 29,000x |
| Voltage (kV) | 300 keV | 300 keV | 300 keV | 300 keV | 300 keV | 300 keV | 300 keV | 300 keV | 300 keV |
| Electron exposure (e–/Å$^2$) | 66 | 66 | 66 | 66 | 66 | 66 | 66 | 66 | 66 |
| Defocus range (μm) | -0.5 to -2.0 | -0.5 to -2.0 | -0.5 to -2.0 | -0.5 to -2.0 | -0.5 to -2.0 | -0.5 to -2.0 | -0.5 to -2.0 | -0.5 to -2.0 | -0.5 to -2.0 |
| Pixel size (Å) | 0.826 | 0.826 | 0.826 | 0.826 | 0.826 | 0.826 | 0.826 | 0.826 | 0.826 |
| Symmetry imposed | C1 | C1 | C1 | C1 | C1 | C1 | C1 | C1 | C1 |
| Initial particle images (no.) | 5,688,448 | 5,688,448 | 5,688,448 | 3,356,580 | 3,356,580 | 3,356,580 | 3,356,580 | 2,031,079 | 2,031,079 |
| Final particle images (no.) | 616,330 | 41,190 | 252,647 | 872,447 | 356,424 | 153,276 | 61,483 | 359,126 | 142,294 |
| Map resolution (Å) | 2.10 | 2.92 | 2.42 | 2.13 | 2.20 | 2.56 | 2.76 | 2.39 | 2.70 |
| FSC threshold | 0.143 | 0.143 | 0.143 | 0.143 | 0.143 | 0.143 | 0.143 | 0.143 | 0.143 |
| **Refinement** | | | | | | | | | |
| Initial model used (PDB code) | Closed state | Open state | 1SXJ | | Closed state | | Open state | Closed state | Open state |
| Model resolution (Å) | 2.03 | 2.90 | 2.30 | | 2.22 | | 2.82 | 2.34 | 2.68 |
| FSC threshold | 0.5 | 0.5 | 0.5 | | 0.5 | | 0.5 | 0.5 | 0.5 |
| Model composition | | | | | | | | | |
| Non-hydrogen atoms | 21,170 | 21,451 | 21,513 | | 23,128 | | 22,779 | 21,7998 | 21,654 |
| Protein residues | 2597 | 2,599 | 2,608 | | 2,758 | | 2,757 | 2,608 | 2,600 |
| Ligands | 9 | 9 | 9 | | 9 | | 9 | 9 | 9 |
| *B* factors (Å$^2$) | | | | | | | | | |
| Protein (mean) | 20.0 | 81.4 | 73.8 | | 59.7 | | 67.0 | 43.1 | 67.0 |
| Ligand (mean) | 11.6 | 47.8 | 41.7 | | 43.6 | | 43.0 | 37.4 | 43.0 |
| R.m.s. deviations | | | | | | | | | |
| Bond lengths (Å) | 0.002 | 0.002 | 0.002 | | 0.002 | | 0.002 | 0.002 | 0.002 |
| Bond angles (°) | 0.460 | 0.530 | 0.517 | | 0.488 | | 0.517 | 0.521 | 0.517 |
| Validation | | | | | | | | | |
| MolProbity score | 1.00 | 1.30 | 1.16 | | 1.14 | | 1.23 | 1.10 | 1.23 |
| Clashscore | 2.24 | 4.76 | 3.75 | | 3.52 | | 4.36 | 3.11 | 4.36 |
| Poor rotamers (%) | 0.74 | 0.00 | 0.95 | | 0.82 | | 0.00 | 0.95 | 0.00 |
| Ramachandran plot | | | | | | | | | |
| Favored (%) | 98.80 | 97.75 | 98.80 | | 98.50 | | 97.95 | 98.73 | 97.95 |
| Allowed (%) | 1.20 | 2.25 | 1.20 | | 1.46 | | 2.05 | 1.27 | 2.05 |
| Disallowed (%) | 0.00 | 0.00 | 0.00 | | 0.04 | | 0.00 | 0.00 | 0.00 |
